# ATF4-Mediated Metabolic Stress Response as a Therapeutic Vulnerability in Chordoma

**DOI:** 10.1101/2025.02.27.640253

**Authors:** Lucia Cottone, James Dunford, Eleanor Calcutt, Vicki Gamble, Filiz Senbabaoglu Aksu, Lorena Ligammari, Giorgia Gaeta, John Christianson, Adrienne M Flanagan, Udo Opperman, Adam P Cribbs

## Abstract

Chordoma, a rare primary bone malignancy, currently lacks effective targeted therapies. Despite surgical resection and adjuvant radiotherapy, prognosis remains poor. Recent preclinical studies have highlighted potential therapeutic targets, including the transcription factor TBXT. However, clinical outcomes associated with therapies targeting TBXT remain underexplored or have been modest, warranting further investigation. In this study, we investigated the therapeutic potential of tRNA synthetase inhibitors in chordoma treatment. Focused compound screening identified distinct chemotypes targeting human glutamyl-prolyl-tRNA synthetase (EPRS) as effective in reducing cell viability in chordoma cell lines through an ATF4-mediated stress response rather than through TBXT regulation. Mechanistically significant upregulation of ATF4 and associated stress response genes was identified with consecutive pro-apoptotic DDIT3-mediated cell death. The prototypic EPRS inhibitor halofuginone demonstrated significant tumour growth inhibition in an *in vivo* patient-derived xenograft model. These results suggest that targeting metabolic stress pathways via ATF4 activation presents a novel therapeutic approach for chordoma, warranting further clinical investigation.

## Introduction

Chordoma is an uncommon malignancy presenting primarily along the spinal axis, exhibiting notochordal differentiation and thought to originate from primitive notochordal remnants^1^. This rare primary bone cancer remains challenging to treat, with the median survival rate for affected patients being approximately seven years^2, 3^. The current standard of care for chordoma involves surgical resection complemented by adjuvant proton or photon radiotherapy. Despite these interventions, the prognosis remains poor.

There is currently no targeted therapy approved for chordoma, however recent preclinical studies have identified several novel therapeutic targets^6^. The efficacy of tyrosine kinase inhibitors has been explored^4, 5^ and although these studies have demonstrated some benefit, the clinical outcomes have been modest (Clinical trial NCT03083678), and there has been limited improvement in patient survival rates. The role of immunotherapy in managing chordoma is an active area of investigation, with early studies suggesting possible therapeutic benefits^6^.

The transcription factor TBXT, essential for notochord development, is epigenetically silenced in the human foetus at approximately 12 weeks of gestation, leading to the regression of the notochord^7^. TBXT is expressed in virtually all chordomas, serving as both a diagnostic marker and a potential therapeutic target^1,8^ Evidence suggests that post-transcriptional modifications of histones also regulate TBXT expression^9^, indicating the potential of epigenetic inhibitors as a promising therapeutic strategy to target selectively TBXT.

In a previous study, we employed a targeted screen of epigenetic-modifying compounds on chordoma cell lines, demonstrating that inhibitors of lysine demethylases (KDM) effectively induce cell death *in vitro* by diminishing TBXT expression^9^. However, these compounds exhibit suboptimal pharmacodynamics, rendering them unsuitable for clinical application, and highlighting the need for further molecular screening.

In this study, we report the findings from an enhanced and broadened drug screening effort for chordoma, incorporating additional epigenetic inhibitors and small molecules, aimed at identifying novel therapeutic targets for this malignancy. We have identified two promising compounds that exhibit potential for chordoma treatment, and we elucidate their mechanisms of action, which involve triggering an ATF stress response in both *in vitro* and *in vivo* models.

## Materials and Methods

### Cell lines

Cell lines UCH-1, UCH-2, UCH-7, UM-Chor1 and MUG-Chor were cultured as previously described^10^. All chordoma cell lines derive from sacral tumours except UM-Chor1, which was generated from a clival chordoma (www.chordomafoundation.org). Briefly, cells were cultured in 4:1 Iscove’s Modified Dulbecco’s Medium (IMDM) (Gibco): RPMI 1640 (Gibco) with 10% fetal bovine serum (FBS) (Cat. F9665, Sigma Aldrich) and 1% Penicillin and Streptomycin (cat. 15070063, Life Technologies). Cell lines were regularly quality controlled for cross contaminating of the cel lines by short-tandem-repeat (STR) analysis, and infection by Mycoplasma.

### Compound screening library

Drug screening was conducted as previously described^9^. Briefly, human chordoma cell lines (U-CH1, U-CH2, U-CH7, MUG-Chor, and UM-Chor1) were seeded at 5,000 cells per well in 96-well plates and incubated overnight. Cells were treated with small-molecule inhibitors or vehicle control (0.1% DMSO) for three to six days (see Supplementary Data 1 for compound details). Cell viability was assessed using Presto Blue Cell Viability Reagent (Thermo Fisher Scientific) and normalised to vehicle controls. Each assay was performed in triplicate, with three independent repetitions. IC50 values were determined using a seven-point dose– response curve, starting at 50 μmol/L, and calculated using Prism version 10.

### Proliferation monitoring assay

Cells were seeded in 96-well plates at a density of 5,000 cells per well in 100 μL of complete culture medium and left to adhere overnight. Prior to seeding, cell viability was assessed using trypan blue exclusion, and only cultures with viability >90% were used. Cell imaging was performed using the Celligo (Revvity), an automated high-content imaging system. The instrument was calibrated according to the manufacturer’s guidelines. Plates were placed in the Celligo, and brightfield images were captured using a 4x objective lens. Images were acquired at multiple time points (0, 24, 48, and 72 hours) to monitor cell proliferation and morphology changes. Images were analysed using Celligo’s integrated software, which utilised automated analysis algorithms to obtain cell counts and confluency measurements by detecting cells based on optimised size, shape, and intensity thresholds for each cell line. The cell counting module quantified the number of cells per well, excluding debris and dead cells through appropriate size and circularity filters. Confluency was measured as the percentage of well area occupied by cells, with adjusted thresholds ensuring accurate cell boundary segmentation. Cell proliferation rates were determined by normalising cell counts at subsequent time points to the initial count, generating growth curves and calculating doubling times using exponential growth models.

### RNA sequencing of chordoma cell lines

Total RNA was extracted from cell lines and tumour samples from treated mice using TRIzol reagent, followed by purification with the Direct-zol RNA Miniprep Kit (Zymo Research), which includes an on-column DNase I digestion step to remove genomic DNA. Poly(A) RNA was isolated from the total RNA using the NEBNext Poly(A) mRNA Magnetic Isolation Module (New England Biolabs). First-strand cDNA libraries were prepared using the NEBNext Ultra Directional RNA Library Prep Kit (New England Biolabs) and sequenced in a 41 bp paired-end configuration on a NextSeq 500 platform (Illumina).

### Illumina sequencing data analysis

Sequencing reads were pseudoaligned to the hg38 reference transcriptome using Kallisto^11^. The quality of the pseudoalignment was assessed with FastQC. Differential gene expression analysis was conducted using the DESeq2 package^12^, with genes considered differentially expressed if the adjusted p-value was < 0.01. Heatmaps of differentially expressed genes were generated using the pheatmap and ggplot2 packages in R.

### Nanopore sequencing

Library preparation was conducted using the cDNA-PCR Barcoding Kit V14 (SQK-PCB114.24, Oxford Nanopore Technologies) according to the manufacturer’s protocol. Briefly, the amplified cDNA was end-repaired and dA-tailed using the NEBNext End Repair / dA-Tailing Module (New England Biolabs). The barcoded adapters provided in the kit were then ligated to the prepared cDNA. The barcoded cDNA was purified using AMPure XP beads (Beckman Coulter) to remove excess adapters and enzymes. The concentration and quality of the barcoded libraries were assessed using the Qubit 4 Fluorometer (Thermo Fisher Scientific) and the Agilent 2100 Bioanalyzer. The samples were then pooled and the library was loaded onto a PromethION 24 device (Oxford Nanopore Technologies) using R10.4.1 flow cells. Sequencing was initiated and run according to the manufacturer’s instructions. Real-time base calling was performed using MinKNOW software (Oxford Nanopore Technologies).

### Nanopore sequencing data analysis

Raw sequencing data were processed using Guppy basecaller (v6.5.7, Oxford Nanopore Technologies) using the high accuracy settings to generate high-quality reads. Subsequent data analysis, including alignment, transcript quantification, and differential expression analysis, was performed using Minimap2^13^ for alignment and DESeq2 for differential expression analysis. Quality control of the sequencing data was assessed using FastQC.

### Western blot and qPCR

Western blot was performed as described previously^10^. Antibodies are listed in **Supplementary Table S1**. All antibodies were blocked in 5% BSA and made up in TBS, with 0.01% tween and 5%BSA. qPCR was performed as described previously^9^ using primers reported in **Supplementary Table S2**.

### Animal experiments

Animal experiments were performed at South Texas Accelerated Research Therapeutics (START) under the following IACUC #: START #09-001. The antitumor activity of halofuginone and palbociclib was tested in a Patient-Derived Xenograft (PDX) model, designated SF8894^16^, in 6/12 weeks old female Athymic Nude (Crl:NU(NCr)-Foxn1nu) immune-deficient mice. Tumour fragments were harvested from host animals and implanted SQ in the right flank into immune-deficient mice, allowed to grow for approximately 20 days and the study initiated at a mean tumour volume of approximately 125-250 mm^3^ upon which the tumour bearing animals were randomized into the different groups. Animals bearing xenografts were excluded if they did not reach sufficient size. Animals were dosed daily with halofuginone (1mg/kg in 10% DMSO, 90% Sterile Saline), Palbociclib (55 mg/kg in 50mM Sodium Lactate, pH4) or no treatment as control. Tumour measurements were performed twice weekly by caliper and no ulcerations were allowed. The tumour endpoint in this experiment was 1000mm^3^ or 42 days. Toxicity was assessed by weight change or changes in eating/feeding or mobility. Mice were weighed twice weekly throughout the course of the drug treatment. Tumour tissue was collected at termination of the study after the animals were euthanized. For freezing and storage in liquid nitrogen, a tumour fragment no greater than 9×9mm was placed in a flash freeze vial on ice and transferred to liquid nitrogen for at least 30 minutes and then placed at -80°C for long-term storage. For FFPE, a 5×5mm core fragment was obtained while on ice (avoiding tumour ends and rim) and placed in 10% formalin fixation Samples were sent in formalin for paraffin-embedding 48 hours after collection.

### Immunohistochemistry for evaluating ki-67 expression

Ki-67 expression was evaluated using immunohistochemistry (IHC). Tissue sections (5 mm) were cut from Formalin fixed, paraffin embedded (FFPE) tissue blocks. Deparaffinization and antigen retrieval were performed using EnVision FLEX Target Retrieval Solution, High pH (DAKO, Agilent technologies) in a PT Link pre-treatment module (DAKO, Agilent). Slides were incubated at room temperature with Ready Probes Endogenous HRP and AP Blocking Solution (Invitrogen) for 15 minutes to block endogenous peroxidase activity. Slides were incubated for a further 20 minutes at room temperature with phosphate-buffered saline (PBS) solution containing 10% normal goat serum (Abcam) and 3% bovine serum albumin (BSA, Sigma-Aldrich) to block non-specific protein binding.

Slides were then incubated with Ki-67 antibody (Polyclonal, ProteinTech - 27309-1-AP) for 30 minutes at room temperature and washed in PBS. Ki-67 signal was detected with diaminobenzidine (DAB) substrate provided with the Mouse and Rabbit Specific HRP/DAB IHC Detection Kit – Micropolymer (Abcam) according to manufacturer’s instruction. Slides were counterstained with haematoxylin (Sigma-Aldrich), sequentially dehydrated with increasing concentrations of ethanol and xylene and mounted using DPX mounting medium (Fisher Scientific). Whole slide images were acquired with Motic EasyScan slide scanner (Motic).

Digital image analysis of immuno-labelled IHC-Ki67 whole-slide images was performed using the open-source image analysis programme QuPath version 0.5.1^14^. Each immune-lablelled section was reviewed to identify and annotate the area of greatest Ki67 positivity, i.e. the proliferative hot-spot. Scans were imported into QuPath and cell detection was performed. Areas of necrosis, tissue folds and artefacts were excluded from analysis, as previously described^14, 15^.

### Statistics

Data were analysed using GraphPad Prism (version 10). Statistical analyses included one-way ANOVA followed by *post hoc* Tukey’s test for multiple comparisons. Results were expressed as mean ± standard deviation (SD) from a minimum of three independent experiments. A p-value <0.05 was considered statistically significant. For animal model data, day 0 tumour volumes were analysed with either ANOVA followed by Dunnett’s test for three or more groups, or Student’s t-test for comparisons involving fewer than three groups.

### Data availability

The raw RNA-sequencing data have been deposited in the GEO database under the accession number GSE275637. The analysis workflows for processing short-read RNA-seq data are available in the GitHub repository https://github.com/cribbslab/cribbslab, utilising the pseudobulk pipeline. Long-read RNA-seq data were processed using the TallyTriN pipeline, accessible via https://github.com/cribbslab/TallyTriN. Subsequently, reads were mapped to features using the featureCounts workflow available in the cribbslab repository.

## Results

### t-RNA synthetase inhibitors can induce cell apoptosis in chordoma cell lines

We previously conducted a comprehensive screening of a library of tool compounds targeting chromatin proteins, including epigenetic readers, writers, and erasers, in chordoma cell lines^9^. In our current study, we expanded these screens to encompass compounds targeting metabolic pathways, kinases, and novel epigenetic inhibitors. MUG-Chor, UM-Chor1, UCH7, UCH2 and UCH1 chordoma cell lines were treated for three and six days in the presence of the compounds and then cell viability was assessed (**Figure 1A-B**, **Supplementary Data 1**). Our findings confirm previous results, demonstrating that histone deacetylase inhibitors, demethylase inhibitors, and PIM kinases are effective in killing three chordoma cell lines (UCH-1, MUG-Chor, and UM-Chor1). Specifically, we reaffirmed dependencies on H3K27me3 demethylase (KDOBA67^9^) and HDAC targets (Belinostat^17^).

**Figure 1.**
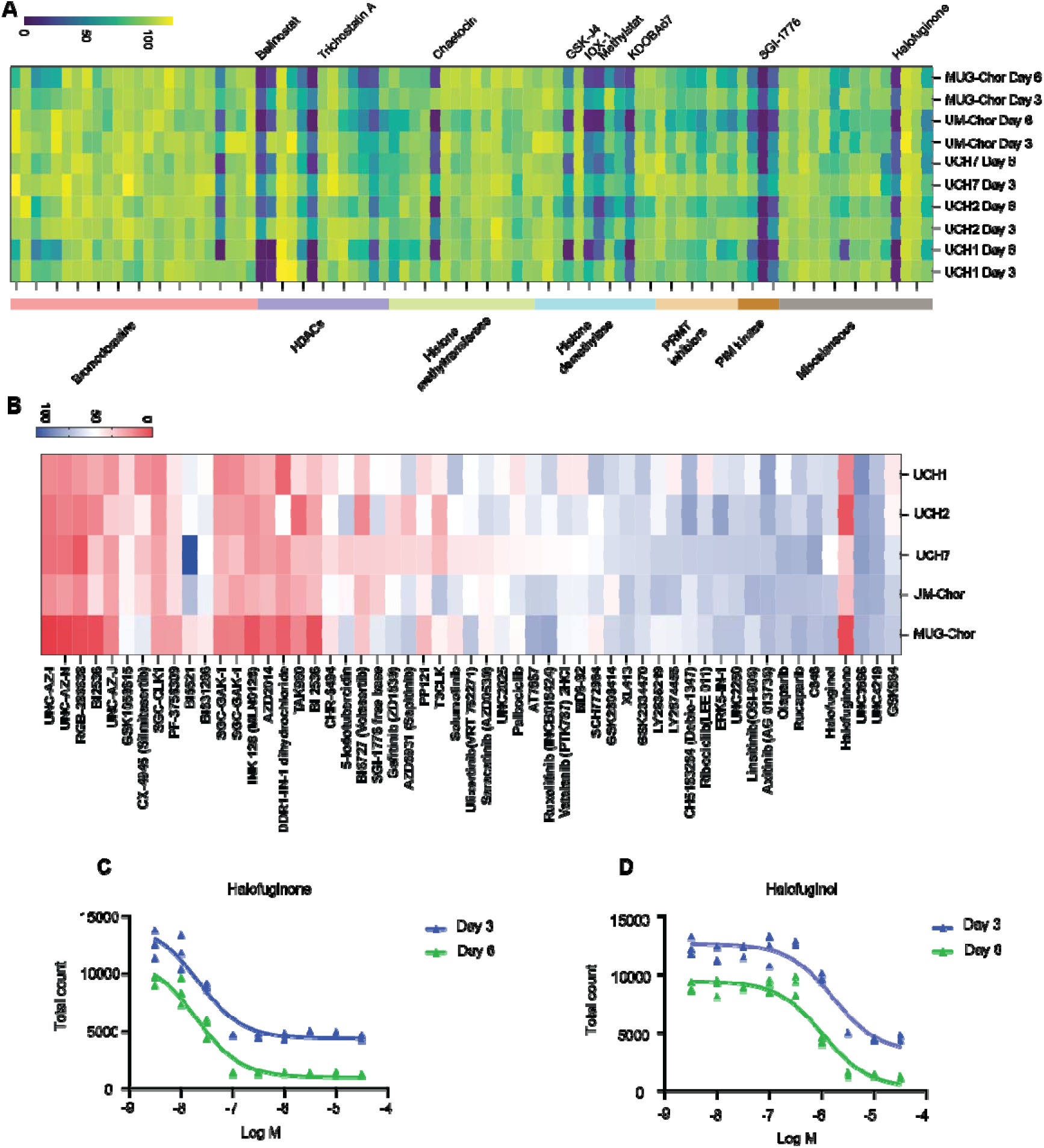
Identification of halofuginone as an Effective Agent in Chordoma through Drug Screening Targeting Metabolic Pathways, Kinases, and Novel Epigenetic Inhibitors. **A-B.** Heatmaps illustrating the screening of 229 small-molecule probes targeting enzymes involved in metabolic pathways, epigenetic modifications (A), and kinases (B) across five chordoma cell lines. Each row corresponds to the mean expression of three replicates, while each column represents an individual compound, grouped by inhibitor class. The data are presented as fractional viability relative to the vehicle (DMSO) control. N=3 replicates per condition. **C-D**. Dose–response curves for halofuginone and halofuginol treatment in the UM-Chor1 chordoma cell line over 3 and 6 days. Each point denotes the mean of three independent experiments. N=3 replicates per condition.

Both screening libraries included the tRNA synthetase inhibitor halofuginone, which reduced cell viability across all cell lines (**Figure 1A**). Halofuginone, a febrifugine quinazoline alkaloid derivative, inhibits the ProRS activity of GluProRS (EPRS) in a proline-competitive manner^18^. Additionally, the library included halofuginol, a tRNA synthetase inhibitor with reduced potency. Our results show that both halofuginone and halofuginol effectively induce cell death in chordoma cell lines, with EC50 values in the low micromolar range (**Figure 1C-D**). EPRS inhibitors have also shown promise in treating multiple myeloma^19, 20^ and SARS-CoV-2^21^ infection.

### Convergent Transcriptional Reprogramming by halofuginone and halofuginol in Chordoma

To investigate the transcriptional changes induced by halofuginone and halofuginol treatments, given their potential to induce apoptosis in chordoma cell lines, we performed RNA sequencing on MUG-Chor cell lines treated with these compounds for three days at their EC50 concentrations (**Figure 2**, **Supplementary Data 2**). Principal component analysis (PCA) of the gene expression data revealed a clear separation between the treated and control groups, with halofuginone and halofuginol treatments clustering closely together, indicating similar transcriptional profiles (**Figure 2A**). This similarity is further supported by the significant overlap in differentially expressed genes between the two treatments (**Figure 2B-D**). Specifically, 82 genes were commonly regulated, and the log2 fold changes in gene expression showed a strong correlation, underscoring the overlapping effects of these compounds (**Figure 2E-F**). Despite these significant gene expression changes, GO and KEGG pathway analyses did not reveal notable pathway enrichments. However, a consistent downregulation of CDC20B (cell division cycle 20B) was observed in both datasets, suggesting a role in cell cycle regulation.

**Figure 2.**
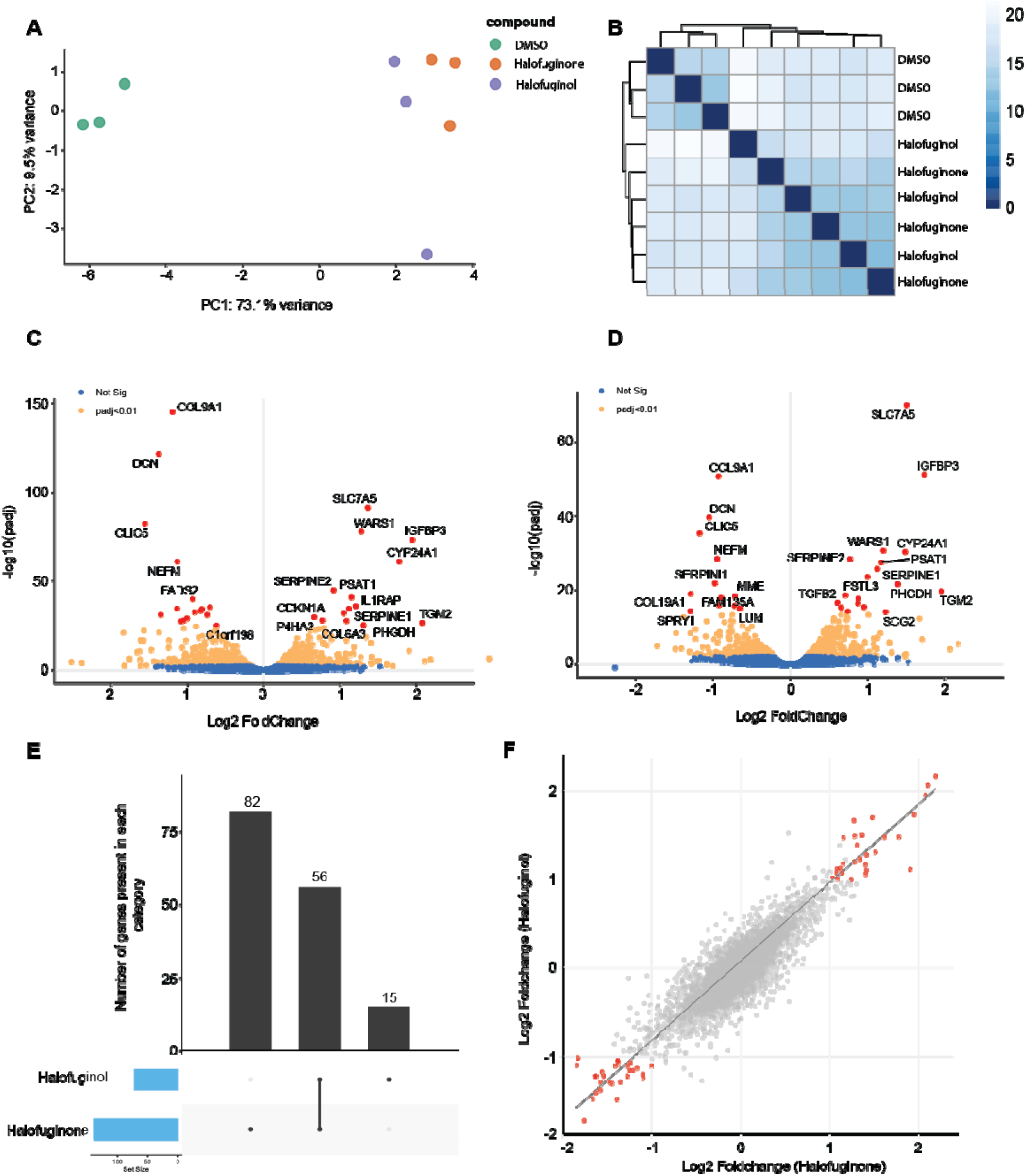
Transcriptomic Profiling of Chordoma Cells in Response to Halofuginone and Halofuginol. **A.** Principal Component Analysis (PCA) of MUG-chor cells treated with DMSO, halofuginone, or halofuginol for 3 days. Each dot represents a biological replicate (n=3 replicates per condition), demonstrating the variance in transcriptomic profiles across treatments. **B.** Heatmap of pairwise Euclidean distances among transcriptomic profiles of MUG-chor cells treated with DMSO, halofuginone, or halofuginol for 3 days, illustrating the degree of similarity or dissimilarity between conditions. **C-D**. Volcano plots depicting differential gene expression in MUG-chor cells treated with halofuginone (**C**) or halofuginol (**D**) for 3 days. Genes with an adjusted P value (Padj) < 0.05 are highlighted in orange, while non-significant genes are shown in blue. Data represent the average of three biological replicates. **E.** Bar plot illustrating the number of differentially regulated genes across treatment conditions: halofuginone alone (left bar), jointly regulated by both halofuginone and halofuginol (middle bar), and halofuginol alone (right bar). **F**. Scatter plot displaying the correlation of Log2 fold changes in gene expression between halofuginone and halofuginol treatments, with each point representing an individual gene.

### Early response genes to tRNA synthetase inhibitors suggest a stress-induced mechanism of action

To elucidate further the mechanisms of action of halofuginone and halofuginol, we investigated early gene responses, prompted by the significant cell death observed by day three of treatment. Transcriptome analysis on MUG-Chor cells was performed at multiple early time points (0, 6, 24, and 72 hours) to capture the dynamic changes in gene expression. Time course analysis of gene expression revealed that the largest source of variance was the treatment duration, followed by the treatment type (**Figure 3A-C**). This analysis highlighted distinct patterns of early and late gene activation. Differentially expressed genes throughout the time course were associated with several key metabolic pathways, including the aerobic electron transport chain, ATP synthesis, and processes regulating DNA damage and protein folding (**Figure 3D**). Importantly, a significant upregulation of stress response genes DDIT3 and ATF4 was observed, with both showing marked increases at the 6hr time points compared to the DMSO control (**Figures 3E-F**). This upregulation persisted, but without significant differential expression between halofuginone and halofuginol, at the 24 and 72-hour time points relative to the DMSO control (**Figures 3E-F**).

**Figure 3.**
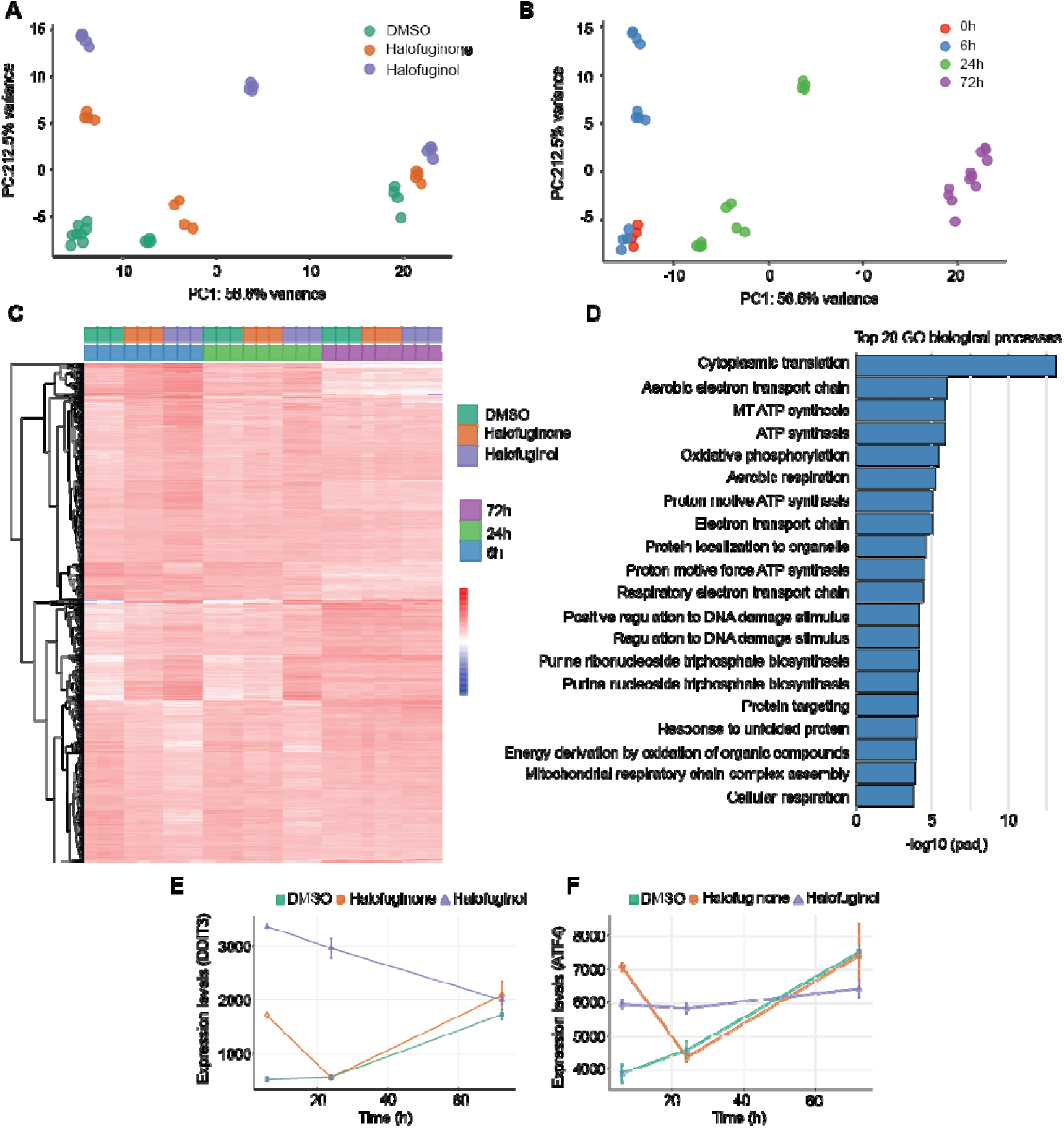
Transcriptomic Analysis of Early and Late Gene Expression Reveals tRNA Synthetase Inhibitors Induce a Stress-Related Mechanism of Action. **A-B.** Principal Component Analysis (PCA) of MUG-chor cells treated with DMSO, halofuginone, or halofuginol at the indicated time points. PCA plots are clustered by treatment (A) or by time point (B). Each dot represents a biological replicate, highlighting the variance in transcriptomic profiles based on treatment and temporal progression. **C.** Heatmap displaying the 1,349 most variably expressed genes in MUG-chor cells treated with DMSO, halofuginone, or halofuginol across the indicated time points. The heatmap illustrates differential gene expression patterns and clustering of genes based on treatment and time course. **D.** Gene Ontology (GO) analysis of differentially expressed genes in halofuginone-treated cells over the full time course (0 to 72 hours). The bar graph ranks the GO terms according to their significance, emphasizing the biological processes most impacted by the treatment. **E-F.** Expression levels of DDIT3 and ATF4 as assessed by RNA-seq analysis in MUG-chor cells over 6hr, 24hr and 72hr following treatment with DMSO, halofuginone, or halofuginol. The line plots represent the dynamic changes in expression levels of these stress response genes, indicating activation of stress pathways. N=3 biological replicates per condition.

### tRNA synthetase inhibition is associated with ATF4 stress response but not with TBXT modulation

We then assessed whether these treatments altered the expression of TBXT, a gene previously reported to be responsive to other compounds^4, 9, 22^. TBXT expression remained unchanged in response to halofuginone nor halofuginol, both at the gene expression and protein level in multiple chordoma cell lines (**Figure 4A-B**). Instead, these compounds induced a pronounced ATF4 stress response, as evidenced by the upregulation of ATF4 targets such as DDIT3 (**Figure 4C**) and the activation of ATF-eIF2α pathways with a reduction in PERK activity and an increase in phosphorylated eIF2a (**Figure 4D**). Therefore, the increase in cell death is not directly related to TBXT but is associated with the activation of an ATF4 stress response.

**Figure 4.**
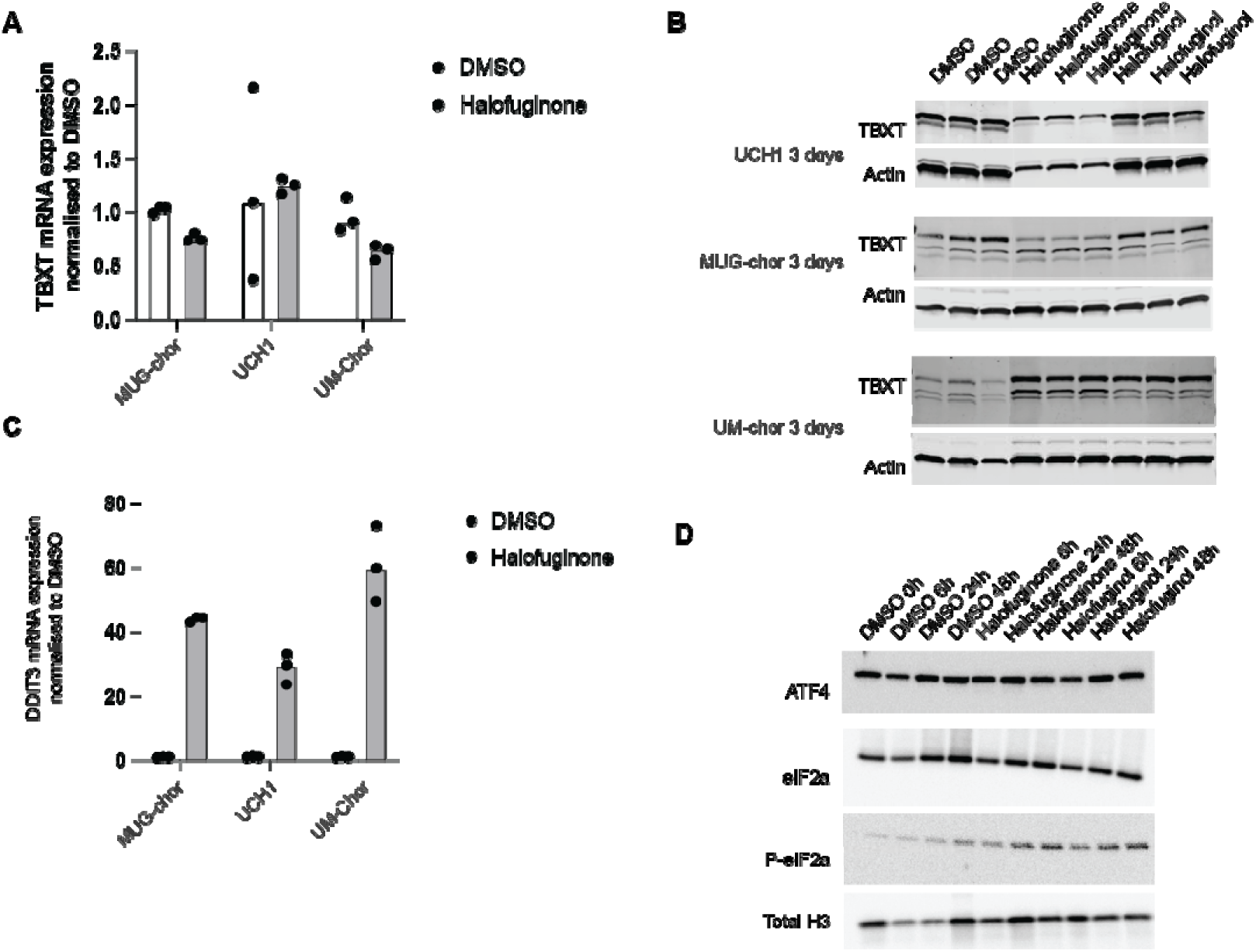
tRNA Synthetase Inhibition Triggers ATF4-Mediated Stress Response without Modulating TBXT Expression. **A.** qPCR analysis of TBXT mRNA expression in chordoma cell lines treated with halofuginone or DMSO. TBXT mRNA expression remains unchanged. Data represent the mean ± standard deviation of three biological replicates (n=3 per condition). **B.** Western blot analysis of TBXT protein expression in multiple chordoma cell lines following three days of treatment with halofuginone or halofuginol. Actin is included as an endogenous loading control. Each condition is represented by three biological replicates. **C.** qPCR analysis of the ATF4 target gene, DDIT3, in chordoma cell lines treated with halofuginone or DMSO. DDIT3 is significantly induced. Data represent the mean ± standard deviation of three biological replicates (n=3 per condition). **D.** Western blot analysis of key proteins in the ATF4-eIF2α stress response pathway, including PERK, ATF4, eIF2α, and phospho-eIF2α (p-eIF2α), in MUG-Chor cells treated with halofuginone or halofuginol at various time points. Total Histone H3 serves as an endogenous control. This panel highlights the activation of the stress response pathway over time.

### Halofuginone reduces chordoma tumour volume in an in vivo xenograft model

Finally, the anti-tumour effects of the compounds were evaluated *in vivo* using the SF8894 PDX model of chordoma (**Supplementary Data 3**). Halofuginone was tested alongside Palbociclib, a kinase inhibitor currently in clinical trials (Clinical Trial ref: NCT03110744), which served as a positive control. Both agents were generally well-tolerated; however, halofuginone treatment resulted in a dose-dependent weight loss of up to 15% at week 40 (**Supplementary Figure 1**). Both Palbociclib and halofuginone demonstrated statistically significant tumour growth inhibition (p<0.05) in the SF8894 model (**Figure 5A**), highlighting the therapeutic potential of halofuginone for chordoma.

**Figure 5.**
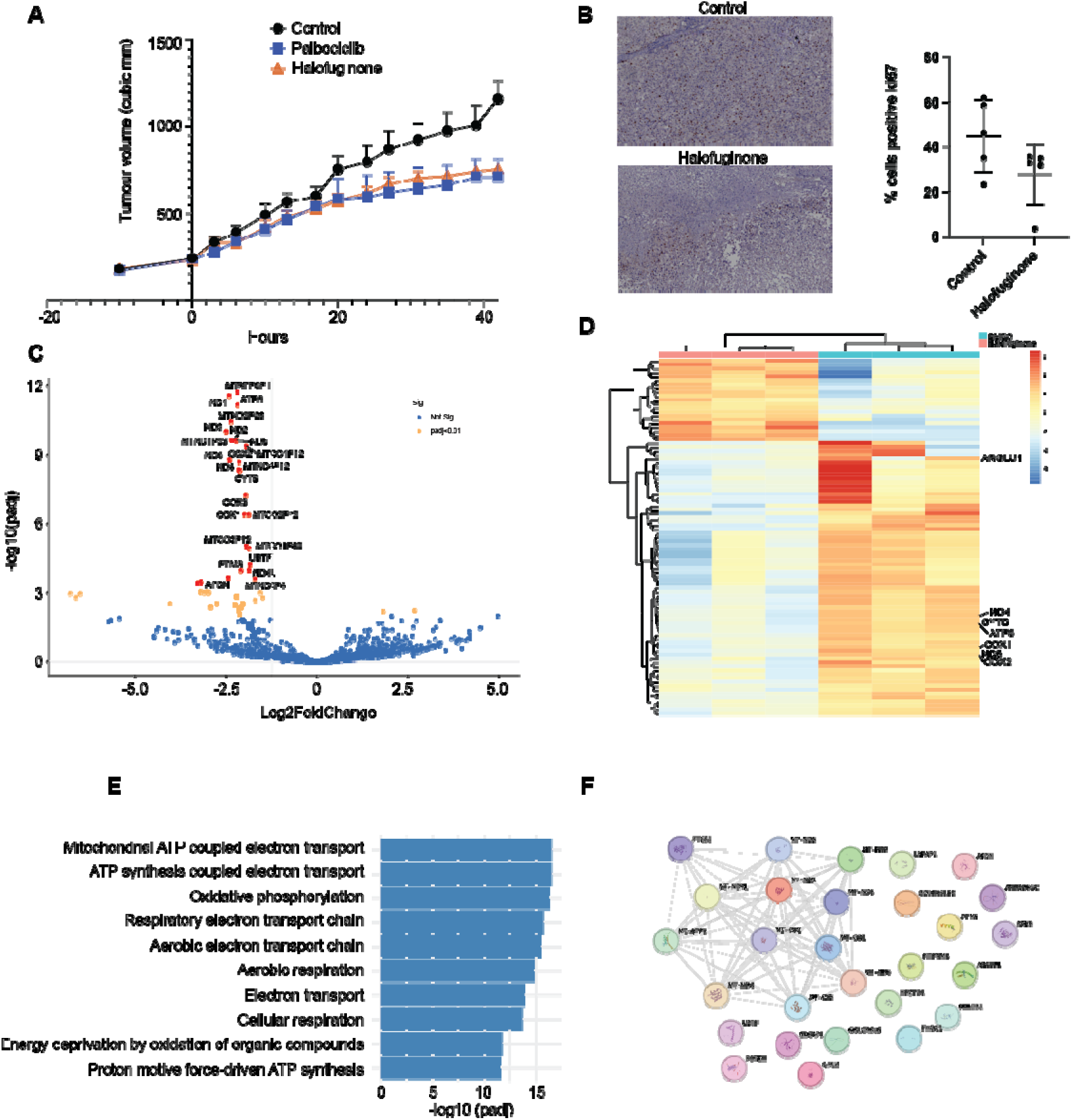
Halofuginone Reduces Chordoma Tumour Volume in an In Vivo Xenograft Model. **A.** Halofuginone treatment results in a significant reduction in tumour growth in the patient-derived xenograft SF8894 mouse model, with effects comparable to the reference standard Palbociclib. Tumour volumes were measured over time, and data are presented as mean ± standard deviation for each treatment group. **B.** Representative immunohistochemistry images (10x magnification) showing Ki-67 staining in tumour tissues from the SF8894 xenograft model after 40 days of treatment with either vehicle control or halofuginone. The right panel quantifies the percentage of Ki-67 positive cells in each group, highlighting a reduction in proliferation in the halofuginone-treated group. **C.** Volcano plot illustrating differential gene expression in tumour tissues from the SF8894 xenograft model following 40 days of halofuginone treatment. Genes with an adjusted P value (Padj) < 0.05 are highlighted in orange, while non-significant genes are shown in blue. Data are based on a minimum of four biological replicates. **D.** Heatmap depicting the most significantly altered genes in tumour tissues from the SF8894 xenograft model after halofuginone treatment. The heatmap illustrates distinct gene expression patterns in response to treatment. **E.** Gene Ontology (GO) analysis of differentially expressed genes in halofuginone-treated tumour tissues, with the top GO terms ranked by significance. **F.** Protein-protein interaction network analysis of significantly altered genes in the halofuginone-treated group, highlighting key pathways and interactions involved in the drug’s mechanism of action.

Immunohistochemical analysis of treated tumours showed a reduction in Ki-67 expression, correlating with the observed decrease in tumour size. We also performed long-read RNA sequencing to conduct full-length transcript analysis on tumour samples from the PDX model treated with DMSO and halofuginone. However, due to sample degradation, reliable full-length transcript analysis could not be achieved, and only gene-level analysis was performed. Despite this limitation, the analysis revealed that most of differentially regulated genes were downregulated in response to halofuginone treatment (**Figure 5B-C** and **Supplementary Data 4**). Gene Ontology and STRING analyses (**Figure 5E-F**) identified an enrichment of genes associated with ATP-dependent pathways, energy deprivation, and significant regulation of mitochondrial respiratory-related genes (**Figure 5E**).

In conclusion, we have identified novel therapeutic candidates for chordoma, demonstrating both *in vitro* and *in vivo* efficacy. These compounds exert their effects through metabolic rewiring of chordoma cells, independent of TBXT.

## Discussion

Our study identifies tRNA synthetase inhibitors, specifically halofuginone and halofuginol, as promising therapeutic agents for chordoma, a malignancy that currently lacks effective targeted treatments^4, 10, 23^. These compounds induced apoptosis in chordoma cell lines through mechanisms independent of TBXT regulation, instead triggering a pronounced ATF4-mediated stress response. The induction of an ATF4 stress response, rather than modulation of TBXT, represents a significant departure from conventional approaches targeting chordoma^24, 25^. Our results align with previous studies highlighting the role of amino acid deprivation and subsequent activation of the integrated stress response (ISR) in immune and cancer cells^20, 26, 27^. Notably, halofuginone and halofuginol treatment resulted in the upregulation of stress response genes such as DDIT3 and ATF4, implicating these pathways in the observed apoptotic effects.

This ATF4-mediated mechanism contrasts with earlier studies on histone deacetylase inhibitors and KDM demethylase inhibitors, which primarily focused on the suppression of TBXT expression^9^. While these earlier studies suggested that TBXT repression could be a viable strategy for chordoma treatment^23^, our findings indicate that targeting metabolic stress pathways may offer an alternative and potentially more effective approach.

In our *in vitro* and *in vivo* model systems halofuginone and halofuginol was effective at reducing chordoma cell viability. Halofuginone significantly inhibited tumour growth, demonstrating its potential as a therapeutic agent for chordoma. This is particularly noteworthy given that halofuginone has already shown promise in treating other conditions, such as multiple myeloma^19, 20^ and SARS-CoV-2 infection^21^, and has received orphan drug status for scleroderma^28^. Our study also highlights the importance of early gene response analysis to understand the mechanisms underlying drug efficacy. Time course analysis revealed distinct early and late gene activation responses, with early upregulation of ATF4 and related stress response genes. This temporal gene expression pattern underscores the dynamic nature of the cellular response to tRNA synthetase inhibition and suggests that early intervention in these stress pathways may be critical for therapeutic success.

In conclusion, our findings demonstrate that tRNA synthetase inhibitors, specifically halofuginone and halofuginol, induce apoptosis in chordoma cells through an ATF4-mediated stress response, independent of TBXT regulation. This mechanism of action provides a new avenue for therapeutic development in chordoma, complementing existing strategies that target epigenetic modifications and transcriptional regulation. Further studies are warranted to optimise these compounds for clinical use and to explore their potential therapeutic benefit in combination with other treatments.

## Acknowledgements and funding

Research support was obtained from Bone Cancer Research Trust, Sarcoma UK (SUKG01.2018), Chordoma UK, Chordoma Foundation. A.P.C. is a recipient of a Medical Research Council career development fellowship (grant no. MR/V010182/1). L.C. and A.M.F. are supported by the University College London Hospitals Biomedical Research Centre, the Cancer Research UK University College London (CRUK-UCL) Experimental Cancer Medicine Centre and the CRUK-UCL Centre Award [C416/A25145]. L.C., a Bone Cancer Research Trust (BCRT) Early Career Fellow, was previously supported by The Tom Prince Cancer Trust. Research in UO laboratory was supported by the Leducq foundation through LEAN program (Leducq Epigenetics in Atherosclerosis Network), and the Chordoma Foundation.

## Competing interests

A.P.C. and U.O. are cofounders of Caeruleus Genomics Ltd and are inventors on several patents related to sequencing technologies filed by Oxford University Innovations.

## Contributions

A.P.C. and U.O designed the study with contributions from L.C. L.C., A.P.C, E.C., V.G, F.S.A, L.L acquired the data. A.P.C. also conducted data analysis. A.P.C. and L.C generated the figures. A.P.C. and L.C. wrote the paper with input from all authors.

## References

1. Sbaraglia, M., Bellan, E. & Dei Tos, A.P. The 2020 WHO Classification of Soft Tissue Tumours: news and perspectives. Pathologica 113, 70–84 (2021).

2. Walcott, B.P. et al. Chordoma: current concepts, management, and future directions. Lancet Oncol 13, e69–76 (2012).

3. Stacchiotti, S., Sommer, J. & Chordoma Global Consensus, G. Building a global consensus approach to chordoma: a position paper from the medical and patient community. Lancet Oncol 16, e71–83 (2015).

4. Magnaghi, P. et al. Afatinib Is a New Therapeutic Approach in Chordoma with a Unique Ability to Target EGFR and Brachyury. Mol Cancer Ther 17, 603–613 (2018).

5. de Castro, C.V. et al. Tyrosine kinase receptor expression in chordomas: phosphorylated AKT correlates inversely with outcome. Hum Pathol 44, 1747–1755 (2013).

6. Freed, D.M., Sommer, J. & Punturi, N. Emerging target discovery and drug repurposing opportunities in chordoma. Front Oncol 12, 1009193 (2022).

7. Risbud, M.V. & Shapiro, I.M. Notochordal cells in the adult intervertebral disc: new perspective on an old question. Crit Rev Eukaryot Gene Expr 21, 29–41 (2011).

8. Vujovic, S. et al. Brachyury, a crucial regulator of notochordal development, is a novel biomarker for chordomas. J Pathol 209, 157–165 (2006).

9. Cottone, L. et al. Inhibition of Histone H3K27 Demethylases Inactivates Brachyury (TBXT) and Promotes Chordoma Cell Death. Cancer Res 80, 4540–4551 (2020).

10. Scheipl, S. et al. EGFR inhibitors identified as a potential treatment for chordoma in a focused compound screen. J Pathol 239, 320–334 (2016).

11. Bray, N.L., Pimentel, H., Melsted, P. & Pachter, L. Near-optimal probabilistic RNA-seq quantification. Nat Biotechnol 34, 525–527 (2016).

12. Love, M.I., Huber, W. & Anders, S. Moderated estimation of fold change and dispersion for RNA-seq data with DESeq2. Genome Biol 15, 550 (2014).

13. Li, H. Minimap2: pairwise alignment for nucleotide sequences. Bioinformatics 34, 3094–3100 (2018).

14. Bankhead, P. et al. QuPath: Open source software for digital pathology image analysis. Sci Rep 7, 16878 (2017).

15. Humphries, M.P. et al. Automated Tumour Recognition and Digital Pathology Scoring Unravels New Role for PD-L1 in Predicting Good Outcome in ER-/HER2+ Breast Cancer. J Oncol 2018, 2937012 (2018).

16. Davies, J.M. et al. Generation of a patient-derived chordoma xenograft and characterization of the phosphoproteome in a recurrent chordoma. J Neurosurg 120, 331–336 (2014).

17. Scheipl, S. et al. Histone deacetylase inhibitors as potential therapeutic approaches for chordoma: an immunohistochemical and functional analysis. J Orthop Res 31, 1999–2005 (2013).

18. Keller, T.L. et al. Halofuginone and other febrifugine derivatives inhibit prolyl-tRNA synthetase. Nat Chem Biol 8, 311–317 (2012).

19. Leiba, M. et al. Halofuginone inhibits multiple myeloma growth in vitro and in vivo and enhances cytotoxicity of conventional and novel agents. Br J Haematol 157, 718–731 (2012).

20. Kurata, K. et al. Prolyl-tRNA synthetase as a novel therapeutic target in multiple myeloma. Blood Cancer J 13, 12 (2023).

21. Sandoval, D.R. et al. The Prolyl-tRNA Synthetase Inhibitor Halofuginone Inhibits SARS-CoV-2 Infection. *bioRxiv*, 2021.2003.2022.436522 (2021).

22. Sharifnia, T. et al. Mapping the landscape of genetic dependencies in chordoma. Nat Commun 14, 1933 (2023).

23. Sharifnia, T. et al. Small-molecule targeting of brachyury transcription factor addiction in chordoma. Nat Med 25, 292–300 (2019).

24. Robinson, H., McFarlane, R.J. & Wakeman, J.A. Brachyury: Strategies for Drugging an Intractable Cancer Therapeutic Target. Trends Cancer 6, 271–273 (2020).

25. Halvorsen, S.C. et al. Transcriptional Profiling Supports the Notochordal Origin of Chordoma and Its Dependence on a TGFB1-TBXT Network. Am J Pathol 193, 532–547 (2023).

26. Nie, A., Sun, B., Fu, Z. & Yu, D. Roles of aminoacyl-tRNA synthetases in immune regulation and immune diseases. Cell Death Dis 10, 901 (2019).

27. Sundrud, M.S. et al. Halofuginone inhibits TH17 cell differentiation by activating the amino acid starvation response. Science 324, 1334–1338 (2009).

28. Pines, M. & Spector, I. Halofuginone - the multifaceted molecule. Molecules 20, 573–594 (2015).

