## Supplementary Data3 for "ATF4-Mediated Metabolic Stress Response as a Therapeutic Vulnerability in Chordoma"

***In Vivo* Study Results with halofuginone in the SF8894 PDX Model**

### 1. Introduction

The antitumor activity of halofuginone and palbociclib was tested in a Patient-Derived Xenograft (PDX) model, designated SF8894, in immune-deficient mice. Data collected from this study included animal weights, observations, and tumor dimensions. This information was used to determine agent tolerability based on weight change and gross physiologic changes; and anticancer activity based on tumor growth inhibition or regression with the data analysis endpoint at Day 42.

### 2. Overall Experimental Design

**Table 2—1: Project Details**

| **Animal Strain:** |  |  | Athymic Nude (Crl:NU(NCr)-Foxn1^nu^) |
| --- | --- | --- | --- |
| **Animal Age / Sex:** |  |  | 6-12 weeks / Female |
| **Selected Models:** |  |  | SF8894 |
| **Administration Volume:** |  |  | 10 mL/kg |
| **Control / ROA:** |  |  | No Treatment |
| **Test Agent(s) / ROA:** |  |  | Palbociclib / PO; Halofuginone / PO |
| **SOC / ROA:** |  |  | None |
| **Study Type:** |  |  | gTGI |

**Additional project details:**

The SF8894 is a PDX model from an adult patient with skull-base conventional chordoma.

gTGI is defined as: Study type where each group is ended once it reaches a designated mean TV or timepoint. Stats based on delta TV vs. Day 0 TV; ≥3 groups = ANOVA + Dunnett’s, <3 groups = Student T-test.

### 3. Materials and Methods

**Table 3—1: Details of Standard Model Systems**

| **Animal Supplier:** | Charles River Laboratories | **Feeding/Drinking Time:** | *Ad Libitum* |
| --- | --- | --- | --- |
| **Acclimatization Period:** | >24 Hours | **Room Temperature:** | 70-74°F |
| **Identification:** | Ear Notch | **Relative Humidity:** | 30-60% |
| **Caging Type:** | Sealsafe® Plus Techniplast, USA | **Light Period:** | 12 Hours |
| **Environment Enrichment:** | Irradiated corncob bedding,  nesting sheets, plastic housing | **Data Capture Method:** | Direct Electronic |
| **Feed Type:** | Teklad 2919 (Irradiated)  19% protein, 9% fat, 4% fiber | **Data Capture Instrument(s):** | Digital Scale / Digital Caliper |
| **Drinking Water:** | Reverse Osmosis,  2ppm Cl_2_ | **Data Storage Method:** | Redundant Cloud Server |

The experiments were performed under the following IACUC #: START #09-001.

Tumor fragments were harvested from host animals and implanted SQ in the right flank into immune-deficient mice, allowed to grow for approximately 20 days and the study initiated at a mean tumor volume of approximately 125-250 mm^3^ upon which the tumor bearing animals were randomized into the different groups. Animals bearing xenografts were excluded if they did not reach the sufficient size. Tumor measurements were performed twice weekly by caliper and no ulcerations were allowed. The tumor endpoint in this experiment was 1000mm^3^ or 42 days. Toxicity was assessed by weight change or changes in eating/feeding or mobility. Mice were weighed twice weekly throughout the course of the drug treatment.

Table 3—2: Test Article and Formulation Data

| Test Agent | Lot Number | Expiry Date | Manufacturer | Vehicle | Formulation(s)  (mg/kg) |
| --- | --- | --- | --- | --- | --- |
| **Palbociclib** | PBC-106 | N/A | LC Labs | 50mM Sodium Lactate, pH4 | 75 |
| **Halofuginone** | S8144 | N/A | Selleckchem | 10% DMSO, 90% Sterile Saline | 1 |

Control animals did not receive treatment

### 4. Detailed Outlines

Table 4—1: Detailed Study Outline

| **-N-**  **mice** | T_X_ | Dose  (mg/kg) | ROA / Schedule | Doses Administered | | Endpoint |
| --- | --- | --- | --- | --- | --- | --- |
|  |  |  |  | Total Dosed | T_X_ Day(s) |  |
| 5 | No Treatment | -- | -- | -- | -- | 42 |
| 5 | Palbociclib | 75 | PO / qd to end | 43 | 0-42 | 42 |
| 5 | Halofuginone | 1 | PO / qd to end | 43 | 0-42 | 42 |

Table 4—2: Detailed Sample Collection

| **Group** | **Tumor** | **# Samples Collected** | **Blood** | | |
| --- | --- | --- | --- | --- | --- |
|  |  |  | **Type** | **Time Point(s)** | **# Samples Collected** |
| No treatment | N | 5 | None | N/A | 0 |
|  | FFPE | 5 |  |  |  |
| Palbociclib | N | 5 | None | N/A | 0 |
|  | FFPE | 5 |  |  |  |
| Halofuginone | N | 5 | None | N/A | 0 |
|  | FFPE | 5 |  |  |  |

Tumor tissue was collected at termination of the study after the animals were euthanized. For freezing and storage in liquid nitrogen, a tumor fragment no greater than 9x9mm was placed in a flash freeze vial on ice and transferred to liquid nitrogen for at least 30 minutes and then placed at -80ºC for long-term storage.  For FFPE, a 5x5mm core fragment was obtained while on ice (avoiding tumor ends and rim) and placed in a properly labeled formalin vial.  Samples were sent in formalin for paraffin-embedding 48 hours after collection.

### 5. Results

Table 5—1: Animal Weight and Agent Tolerability

| Group | Weight Data (Day 42) | | Weight Nadir (Day 42) | | Drug Deaths | | | | | | |  |  |
| --- | --- | --- | --- | --- | --- | --- | --- | --- | --- | --- | --- | --- | --- |
|  | **Mean ± SD** | **%vD_0_** | **%vD_0(max)_** | **Day** | **D** | Day | **T** | Day | **B** | Day | **U** | | Day |
| No treatment | 29 ± 2 | +10% | -3% | 3 | **0** | -- | **0** | -- | **0** | -- | **0** | | -- |
| Palbociclib | 25 ± 2 | +3% | -5% | 6 | **0** | -- | **0** | -- | **0** | -- | **0** | | -- |
| Halofuginone | 21 ± 1 | -15% | -15% | 42 | **0** | -- | **0** | -- | **0** | -- | **0** | | -- |

Abbreviations: %vD_0_= Weight change versus a study initiation (Day 0) measurement; %vD_0(max)_= Maximum weight loss versus a study initiation (Day 0) measurement; D= Death as a result of agent toxicity; T= Death as a result of technician error; B= Death as a result of tumor-related weight loss or cachexia; U= Cause of death cannot be determined

Table 5—2: Agent Efficacy and Tumor Volume Data

| Group | Mean ± SEM  (Day 42) | %TGI | p-value | Significant | PR (%TR) | CR | TFS |
| --- | --- | --- | --- | --- | --- | --- | --- |
| No treatment | 1161 ± 103 | -- | -- | -- | 0 | 0 | 0 |
| Palbociclib | 710 ± 105 | 49% | 0.0019 | Y | 0 | 0 | 0 |
| Halofuginone | 755 ± 20 | 44% | 0.0052 | Y | 0 | 0 | 0 |

%TGI is defined as: Percent mean tumor growth at Day 42 versus Day 0 between treatment (TX) and untreated groups; Formula: %TGI = 1 – (TXf_avg_ – TXi_avg_) / (Cf_avg_ – Ci_avg_) where TX=mean tumor growth in drug treated animals; C=mean tumor growth in untreated animals; f=measurement on day 42; i= measurement on day 0. P-values <0.05 were considered statistically significant.

### 6. Summary

Halofuginone was tested in the SF8894 patient derived xenograft model. Results with the CDK4/6 inhibitor, Palbociclib, were included for comparison. Both agents were tolerated with some treatment-dependent weight loss which reached -15% in the halofuginone treated animals. Both palbociclib and halofuginone reported statistically significant (p<0.05 or better) modest tumor growth inhibition towards SF8894.
