## Supplementary Tables for "ATF4-Mediated Metabolic Stress Response as a Therapeutic Vulnerability in Chordoma"

**Supplementary Table S1. List of Antibodies used in the study.**

| **Protein** | **Brand** | **Cat. Number** |
| --- | --- | --- |
| Beta-Actin | Sigma | A5441 |
| TBXT (Brachyury A4) | Santa Cruz | Sc-374321 |
| PERK | Cell Signalling | cat#5683 |
| ATF4 | Cell Signalling | cat#11815 |
| eIF2-alpha | Abcam | cat #ab264253 |
| phospho-eIF2-alpha | Cell signalling | cat #ab32157 |
| Total H3 | Abcam | cat#ab1791 |

**Supplementary Table S2. List of primers used in the study.**

| Gene | Application | Fw Primer | Rev Primer |
| --- | --- | --- | --- |
| TBXT | qPCR | CCCGTCTCCTTCAGCAAAGTC | TGGATTCGAGGCTCATACTTATGC |
| DDIT3 | qPCR | ATGAACGGCTCAAGCAGGAA | GCAGATTCACCATTCGGTCAA |
