## Supplementary figures and images for "ATF4-Mediated Metabolic Stress Response as a Therapeutic Vulnerability in Chordoma"

### Supplementary Figure 1

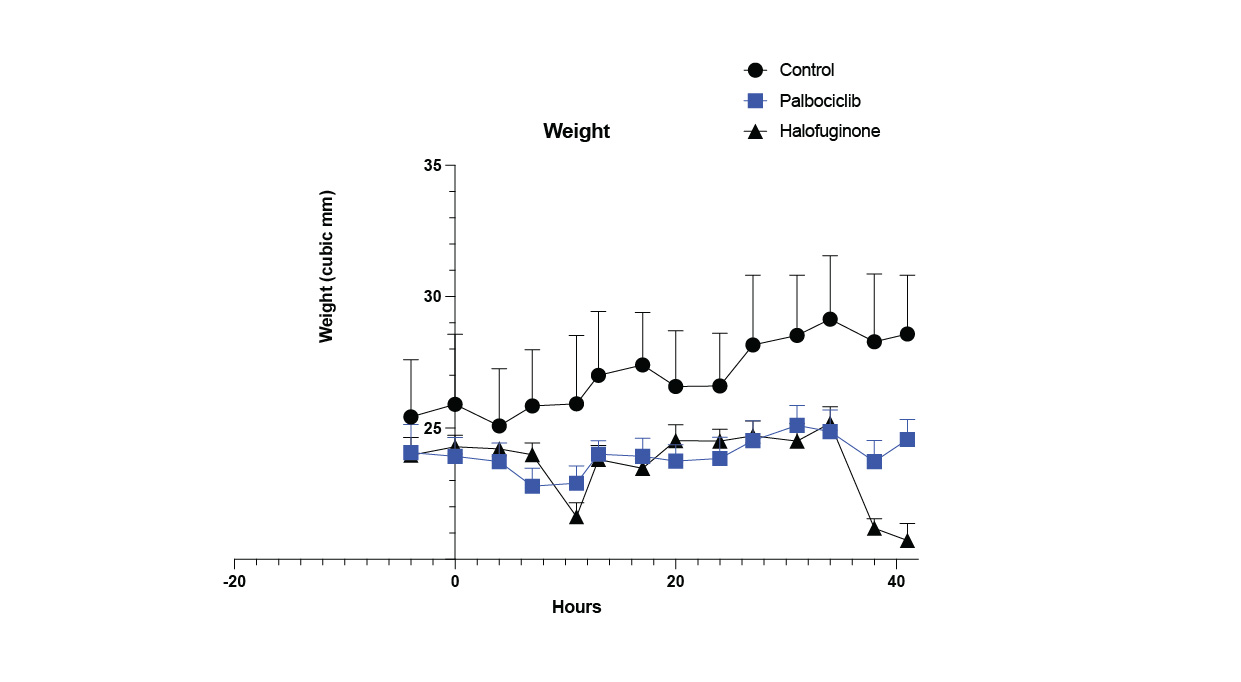
